# Gctf: real-time CTF determination and correction

**DOI:** 10.1101/022376

**Authors:** Kai Zhang

## Abstract

Accurate estimation of the contrast transfer function (CTF) is critical for a near-atomic resolution cryo electron microscopy (cryoEM) reconstruction. Here, I present a GPU-accelerated computer program, Gctf, for accurate and robust, real-time CTF determination. Similar to alternative programs, the main target of Gctf is to maximize the cross-correlation of a simulated CTF with the power spectra of observed micrographs after background reduction. However, novel approaches in Gctf improve both speed and accuracy. In addition to GPU acceleration, a fast ‘1-dimensional search plus 2-dimensional refinement (1S2R)’ procedure significantly speeds up Gctf. Based on the global CTF determination, the local defocus for each particle and for single frames of movies is accurately refined, which improves CTF parameters of all particles for subsequent image processing. Novel diagnosis method using equiphase averaging(EFA) and self-consistency verification procedures have also been implemented in the program for practical use, especially for aims of near-atomic reconstruction. Gctf is an independent program and the outputs can be easily imported into other cryoEM software such as Relion and Frealign. The results from several representative datasets are shown and discussed in this paper.

## 1. Introduction

Recent progress has allowed cryo-electron microscopy (cryoEM) to determine structures of bio-macromolecules to near-atomic resolution(Nogales and Scheres, 2015). This is due to developments in multiple fields, but especially better detectors and image processing methods(Bai et al., 2015). The significantly improved detective quantum efficiency (DQE) of direct detectors, such as Falcon II and K2 summit, makes the quality of the cryoEM reconstructions much better than when using traditional CCD or film(Bai et al., 2015). Recording movies on these detectors allows motion correction of entire micrographs or individual particles, which makes critical improvements for high resolution reconstruction. More and more structures at near-atomic or atomic resolution are being solved recently by cryoEM(Amunts et al., 2015; Bartesaghi et al., 2015; Jiang et al., 2015; Paulsen et al., 2015; Taylor et al., 2015; Urnavicius et al., 2015; Zhao et al., 2015).

In contrast to a simple projection of a 3-dimensional object, the cryoEM image of vitrified specimen is modulated by contrast transfer function (CTF) in Fourier space. Because of the thin vitreous ice film, the image formation can be well described by weak-phase approximation(Wade, 1992). Based on this approximation, the phase contrast is dominant while the amplitude contrast is very small. Therefore, the major factors that affect the CTF of cryoEM image formation are the defocus and aberration of lens. The effect of these factors makes CTF a frequency-dependent oscillatory function, modulating both the amplitudes and phases of the image. Original information of the images must be restored by CTF correction in order to obtain the correct 3D reconstruction. The oscillation of the CTF becomes more severe at higher frequency or under a higher defocus. For this reason, image restoration is quite challenging, especially for the high frequency information, which makes accurate CTF determination an important factor for near-atomic 3D reconstructions.

There are currently several programs available for CTF determination(Ludtke et al., 1999; Mallick et al., 2005; Mindell and Grigorieff, 2003; Penczek et al., 2014; Shaikh et al., 2008; Sorzano et al., 2004; Vargas et al., 2013; Voortman et al., 2011). In a recent work, researchers systematically studied the performance of different programs(Marabini et al., 2015). Each of the programs has its own advantages for certain purposes. The popular program CTFFIND3(Mindell and Grigorieff, 2003) shows the best results using real datasets in this benchmark test, in spite of a slightly lower rank using simulated micrographs. However, with the fast development of cryoEM, a lot of new challenges are being required for daily image processing. One challenging requirement is to further improve CTF accuracy for 3D reconstruction at near-atomic or real atomic resolution. Higher speed without sacrificing the accuracy is also helpful to facilitate data processing with the development of automatic data collection at higher throughput. Besides, automatic self-consistency verification of the CTF determination and quality evaluation of the micrographs will greatly facilitate the ultimate goal of automation in cryoEM.

Here I present a robust GPU-accelerated computer program called Gctf for CTF determination, refinement and correction. GPU acceleration as well as an optimized programming strategy makes Gctf very fast. It can easily process thousands of micrographs within minutes using a single GPU card. The accuracy of the global CTF determination was verified by both manual scrutiny and automatic verification. Astigmatism-based rotational averaging, or what I call Equiphase Averaging (EFA) makes the power spectrum significantly improved for better diagnosis. Gctf was tested using a variety of parameters, showing stable ranges of parameter selection and thus its potential power for CTF automation of many types of micrographs. Micrographs from many datasets collected at the MRC-LMB (Cambridge) and several other collaborating institutes proved the accuracy, speed, convenience and robustness in practical use. In almost all cases, there was no need for parameter optimization. Gctf also performed well with a number of deliberately selected challenging micrographs.

Local refinement and movie processing have also been implemented in Gctf. Local defocus refinement for each single particle makes significant improvements for 3D reconstructions carried out with datasets that have large defocus variation. Refinement of defocus of each frame in a movie provides a way of tracking frame movement in the Z-direction during imaging. Beside the determination and refinement of defocus in Gctf, automatic self-consistency verification and micrographs quality evaluation is also available for better automation of cryoEM data processing.

## 2. Theory and methods

### 2.1 Definition of Contrast transfer function

Image formation in a weak-phase approximation is modulated by the CTF which can be defined as Eq. (1).

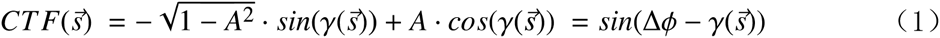

Where 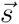 is the spatial frequency; ***A*** is the amplitude contrast coefficient; 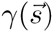 is a function of 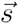 representing the varying phases of the CTF, while Δ*ϕ* is a global phase shift contributed by amplitude contrast. Ideally an image can be regarded as a projection of a 3D object convoluted by the CTF. In other words, the Fourier transform of an image is the Fourier transform of a projection multiplied by the CTF. Note that an envelope function and noise severely affect the real image formation which must be taken into consideration for reliable CTF determination and correction.

Defocus and spherical aberration of the microscope lens are the two major factors that affect the values of 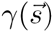 formulated as Eq. (2). The effect by other factors such as coma aberration is ignored in the current CTF determination method.

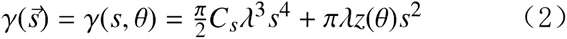

Where *s* is the modulus of 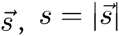, and 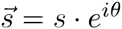; *λ* is the wavelength of an electron; *C*_*s*_ is the spherical aberration coefficient; *z*(*θ*) is the defocus in the direction with an azimuthal angle *θ*, which can be precisely calculated using an elliptic function based on the averaged defocus with astigmatism Eq. (3).

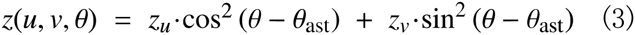

Where the defocus is regarded as a ternary variable *z*(*u*, *v, θ*), *u* and *v* represents the maximum or minimum defocus; *θ* is the varying azimuthal angle; *θ*_*ast*_ is the fixed angle between axis *z*_*u*_ and x-axis of Cartesian coordinate system.

### 2.2 Gctf target

The target of Gctf is trying to maximize the cross-correlation of the simulated CTF amplitudes with the amplitudes of a Fourier transform of the raw micrograph after background reduction. The estimation of the background uses a box-convolution of the natural logarithm of amplitudes ***ln*|*F*|** for estimation of background. In contrast CTFFIND3 uses the original amplitudes of the Fourier transform. I chose ***ln*|*F*|** because it down weights the strong signal at low frequency that tends to dominate and may mislead the fitting. Gctf also uses a B-factor to decrease the simulated CTF at higher frequency to reduce over-fitting of noise. The final target in Gctf is to estimate the defocus *z*(***u***, ***v, θ***), which is the only unknown parameter of the CTF. This estimation is described by Eq. (4)

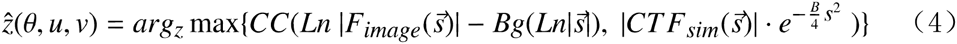

Where, 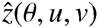 is the estimated CTF parameter; ***F***_*image*_ is the Fourier transform of real data; ***Bg*** is the estimated background; ***CTF***_*sim*_ is the simulated CTF; ***CC*** represents the cross-correlation; ***B*** is B-factor used to down-weight high-frequency. Note that Gctf only normalizes the cross-correlation coefficient at last step for faster speed.

### 2.3 Flow-chart of Gctf

The overall flow chart of Gctf can be described as shown in Figure 1. The preparation step contains the following process: handling input/output parameters; setting up the program running environment (e.g. checking and assigning the GPU device); allocating necessary memory for both CPU and GPU; pre-calculating sharable parameters and data.

**Figure 1.**
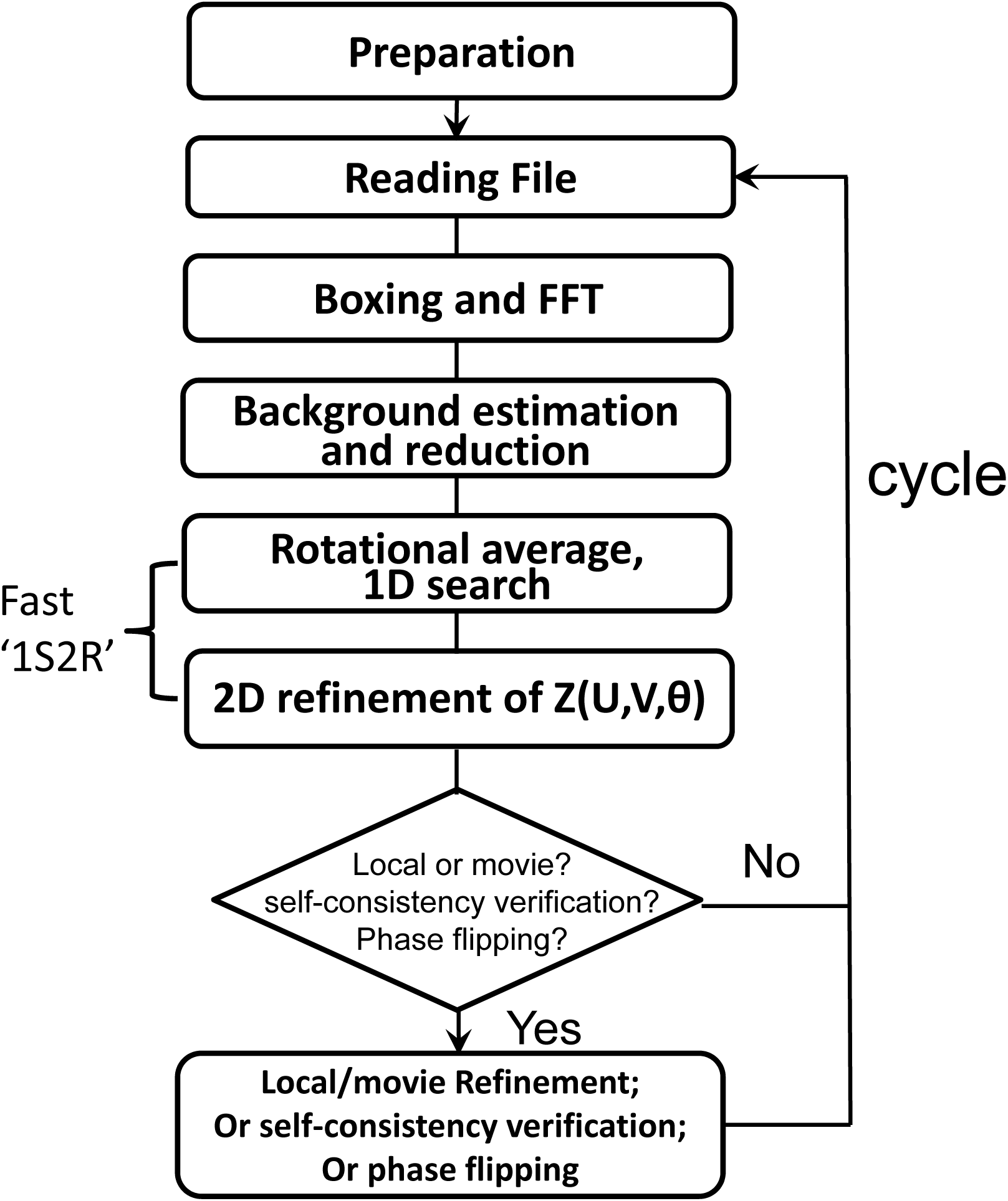
Flow chart of Gctf.

The CTF determination contains the following steps (Figure 1): read and write files; box out sub-areas and perform a series of Fast Fourier Transform (FFT) to generate an averaged power spectrum; estimate and subtract the background; rotationally average the power spectrum to get a 1D profile (Figure 2A); search for the average defocus that best fits the observed 1D profile; perform a 2-dimensional (2D) refinement of all three parameters of defocus *z*(***u***, ***v, θ***). The key procedure of ‘1D search plus 2D refinement’ is called ‘1S2R’ briefly in the rest parts of this paper. This procedure has been proved to be reliable and very fast using practical data(details in Section 3.2 and 3.3). In addition, local and movie refinement, self-consistency verification or phase flipping can be performed if specified. Gctf then reads and processes another file until all have been processed.

**Figure 2.**
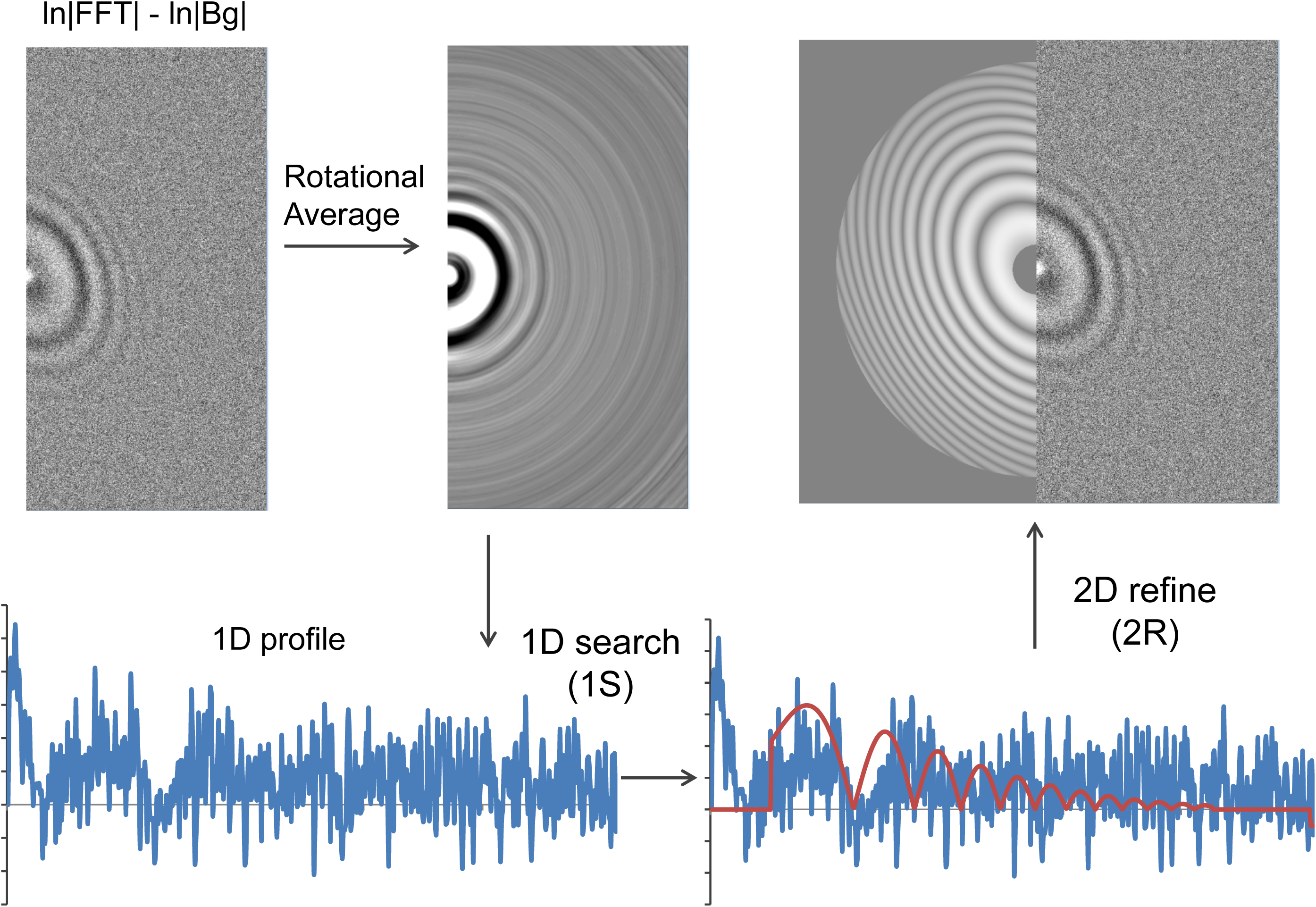
Flow chart of Gctf using a real micrograph. A micrograph with significant astigmatism in dataset-1 is presented to demonstrate the procedure clearly.

The following several sections will describe point-by-point more details of the methods and theories that are used in Gctf.

### 2.4 Defocus inaccuracy related phase error criterion

The accuracy of defocus determination is very important for high-resolution cryoEM reconstructions. It is not easy to calculate the exact phase error between the simulated and observed CTF. However the CTF phase error caused by inaccuracy of defocus estimation can be well predicted. Assuming the difference between the true defocus of a micrograph and the estimated defocus is Δ*z* which causes the phase error Δ*γ*(s).

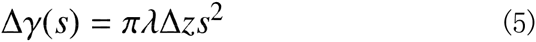

It is clear from the Eq. (5) that the defocus-inaccuracy dependent phase error is proportional to frequency squared for a certain micrograph Eq. (6).

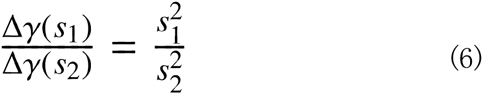

Obviously from Eq. (5) or (6), an error in CTF determination, which can be ignored for a lower resolution reconstruction, might cause a critical error at high resolution. Theoretically, if the phase error is smaller than 90 degrees, there is still information and the defocus value is usable. Based on the 90 degree criterion, I calculated and plotted CTF phase error versus frequency for different defocus errors between 10nm and 200nm (Figure 3a). The maximum allowed CTF defocus errors were plotted against frequency for three typical voltages used in cryoEM reconstructions (Figure 3b). At 300kV, the voltage used for most high resolution reconstructions, the maximum allowed defocus error is 40nm for a 4.0Å resolution reconstruction. For a ˜2.0Å resolution reconstruction, the target defocus accuracy must be better than 10nm. In practice, defocus inaccuracy is only one of the factors that cause CTF phase error. Magnification distortion, chromatic or comatic aberration, astigmatism inaccuracy, as well as mechanical and beam induced movement of the sample can all contribute to the phase error during an experiment. Data processing can also lead to large phase errors, especially at high frequency. Due to these many sources of error, a 90 degree criterion on CTF determination is not good enough to select high quality micrographs.

**Figure 3.**
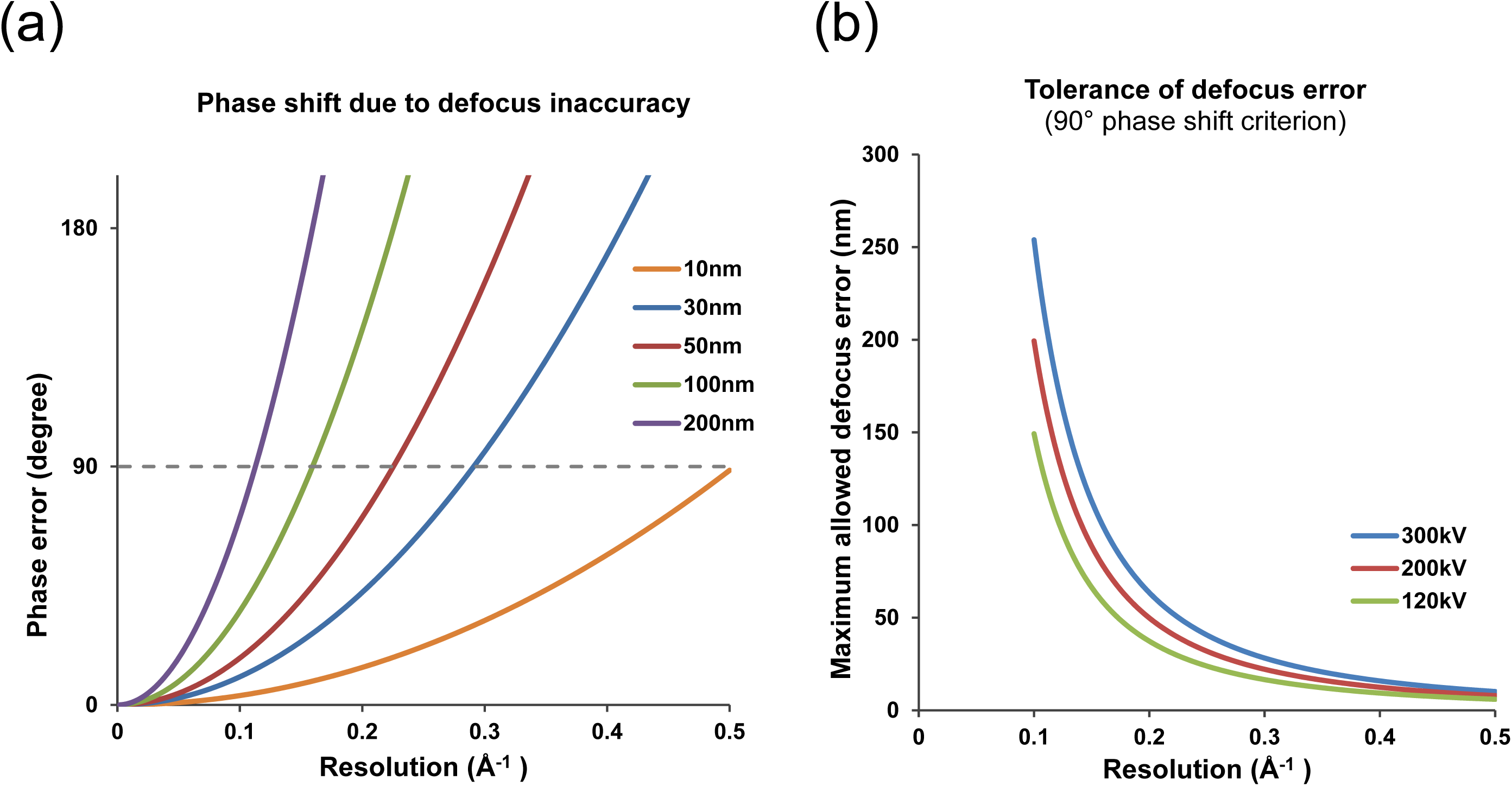
Relationship between CTF phase error and defocus inaccuracy. (a) The errors of CTF phases by different levels of defocus inaccuracy at 300kV high tension. The dashed gray line represents the threshold for 90°phase shift criterion. (b) Based on the 90°criterion from (a), the maximum defocus inaccuracy allowed at various resolutions for three typical high tension values (300kV, 200kV and 100kV) are plotted.

### 2.5 Local refinement strategy

The accuracy of defocus for near-atomic resolution (<4.0Å) should be at least better than 40nm as described above. However, stage tilt, uneven ice, a distorted supporting carbon film or charging can all lead to the defocus variation among particles within a cryoEM micrograph. Simply considering the tilt of micrograph will not generate accurate local defocus caused by nonlinear factors. Therefore, a new local refinement strategy for each particle in one micrograph is implemented in Gctf to solve this problem without assuming any model for defocus variation. Its goal is to refine the defocus of each particle as accurately as possible while minimizing the over-fitting of noise.

Because of the low signal to noise ratio (SNR), direct CTF determination for each particle is not always practical. To best handle this issue, Gctf does a two-step estimation of single particle CTF determination. First, it determines the global CTF parameters for an entire micrograph. Then based on these global values, it does a local refinement for each particle. The target is to estimate the power spectrum of each particle together with its surrounding areas. It uses Gaussian weighting according to the distances between the centers of the particles as described in Eq. (7).

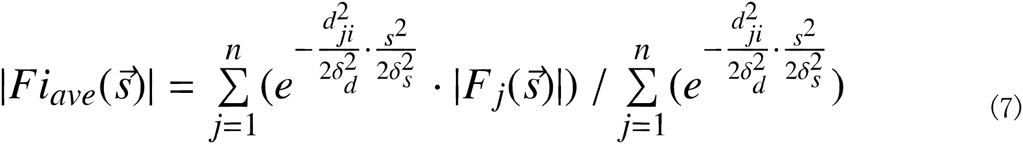

Where |***Fi***_*ave*_**|** is the averaged amplitudes of ***i*_*th*_** particle; |***F*_*j*_|** the amplitudes of ***j_th_*** neighbor; *d*_*ji*_is the distance between particle ***i*** and its neighbor ***j*** and *δ*_*d*_ is the standard deviation of all distances to all neighbors; *δ*_*s*_ is similar to but *δ*_*d*_ with a down-weighting of high-frequency. Note that the combination of the weighting by distance and frequency is a multiplication of the exponent.

There are two different approaches in Gctf for local refinement. One approach simply takes everything in the neighboring areas into account. The other approach uses the coordinates of picked particles or user defined boxes. These two approaches might be useful in different situations. For example, in the case of vitreous ice on a thin carbon layer, using all the neighboring areas will improve the SNR due to the contribution of carbon to the final power spectrum. In the case of pure ice, using neighboring coordinates of picked particles might be better because the pure ice does not contribute as much to the averaged power spectrum.

### 2.6 CTF refinement for movies

One of the biggest advances in cryoEM recently is the invention of direct electron detectors which allow movie recording. Beam induced movement correction using movies has greatly improved the resolution of the final reconstruction(Bai et al., 2013; Li et al., 2013). The movement in the X or Y direction of a micrograph is usually around several Ångstroms, while the Z-direction movement can be over a hundred Ångstroms(Russo and Passmore, 2014). Although the movement is dominantly in the Z-direction, the small movement in the XY plane severely affects the quality of cryoEM micrographs. Motion correction programs normally consider only the drift in the XY plane because the eucentric height of the object does not affect its 2D projection. However, EM micrographs are modulated by CTF, which is sensitive to Z-height changes. Beam induced movement might change the CTF from frame to frame. A hundred Ångstrom movement is not a significant change even up to a 3Å reconstruction, but Figure 3 suggests it might help to improve a reconstruction close to 2Å.

I implemented accurate defocus refinement for movie frames in Gctf to deal with large movement in the Z-direction or defocus changes due to charging. Similar to local defocus refinement, movie defocus refinement is performed in two steps. First, global CTF parameters are determined for the averaged micrograph of motion-corrected movies. Then based on the global values, parameters for each frame are refined, using an average of 5 to 10 adjacent frames to reduce the noise.

### 2.7 Resolution-extention and Bfactor-switch

Strong structure factors at low spatial frequencies can lead to CTF determination bias. Direct CTF determination at high frequency using the ‘1S2R’ procedure might fail in the case of large astigmatism due to severe oscillation of CTF. I provide two options to deal with micrographs that have very large astigmatism. They both make the ‘1S2R’ procedure more robust in such challenging case. One option is ‘resolution-extension(RE)’ and the other is ‘Bfactor-switch(BS)’. In the first method, Gctf determines an initial CTF parameters using a relatively lower resolution shell(e.g. 50–10Å by default). Based on this value, the resolution is then extended to a higher range(e.g. 15 to 4Å) for CTF refinement. In the second method, Gctf uses a larger Bfactor (e.g. 500Å ^2^) to significantly down-weight high frequency for initial CTF determination. Then it switches to a smaller Bfactor (e.g. 50Å ^2^) to refine the previously determined CTF parameters. Either method shows its power to deal with some challenging cases(detailed results in Section 3.5). The combination(REBS) can even work slightly better in certain cases.

### 2.8 Equiphase average (EFA)

The astigmatism of practical datasets can range from several hundred to over a thousand Ångstroms. One of the tested datasets (hepatitis A virus, HAV) had an astigmatism of ˜1800Å but still reached 3.4Å resolution. High astigmatism makes the Thon rings in the power spectrum elliptical, which means simple rotational averaging will not provide a good estimation for them. I therefore use the approach called ‘Equiphase Average (EFA)’. The idea is to average the amplitudes of the micrograph FFT which have the same CTF phases 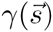 (Figure 4 and Eq. 8).

**Figure 4.**
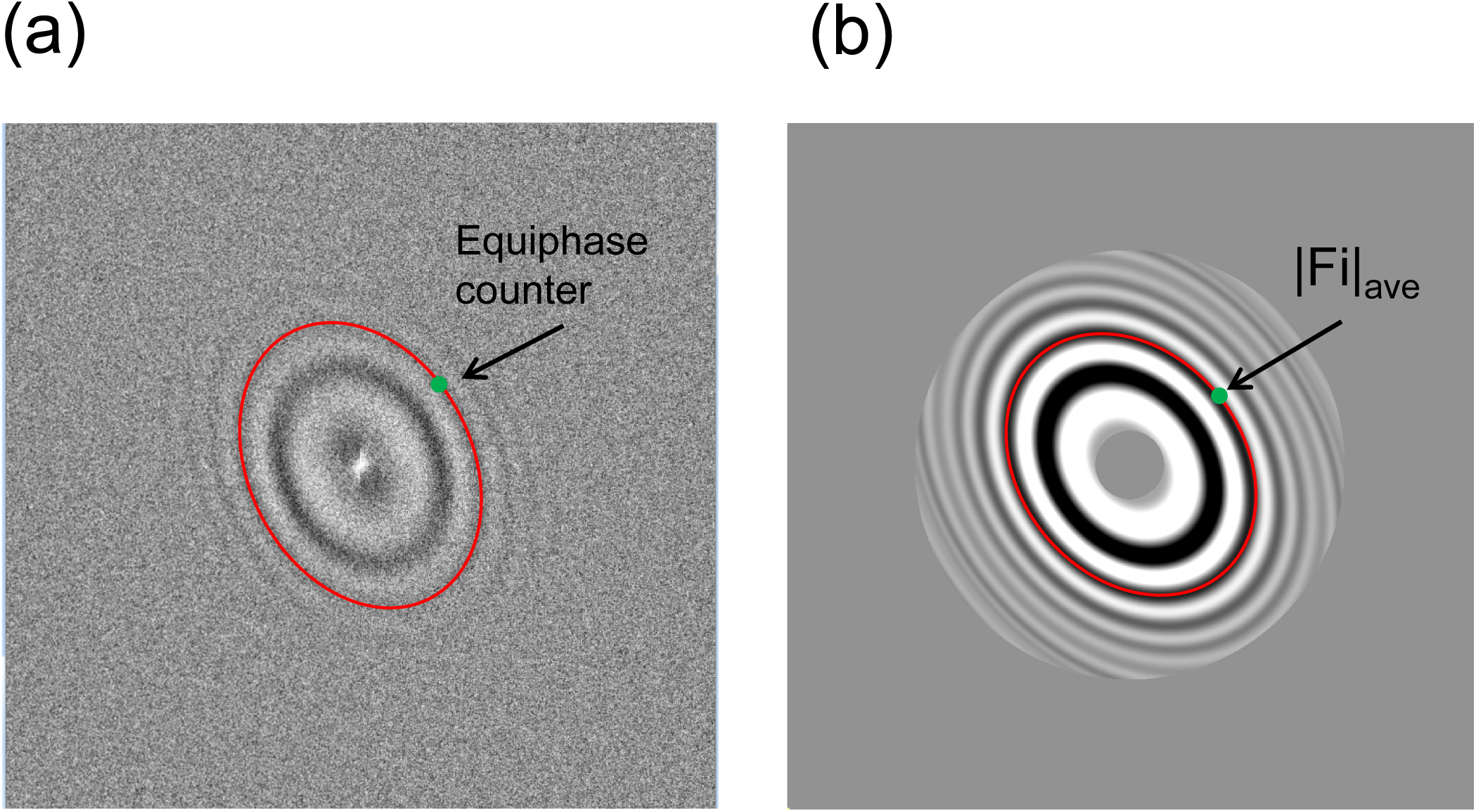
Equiphase average. (a) The logarithmic power spectrum after background reduction. The green point is the target pixel to be averaged. The red line represents all pixels with equiphases for the green point in this image. (b) A typical equiphase averaged power spectrum. Resolution lower than 50Å or higher than 7Å has been excluded.

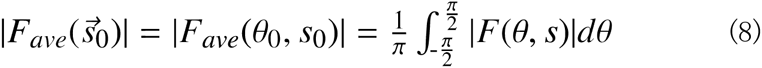

For a specific point with frequency magnitude *s*_0_ and azimuthal angle *θ*_0_ in Fourier space, Eq. (8) represents how the rotational average of the amplitude |***F***_*ave*_**|** is calculated. If the frequency ***s*** is independent of *θ*, or in other words is always equal to *s*_0_, it is the normal rotational average. In the EFA, Gctf only averages the amplitudes with the same CTF phases as in Eq. (9). For any angle *θ*, the frequency magnitude *s*(*θ*) involved in the average is calculated using Eq. (10).

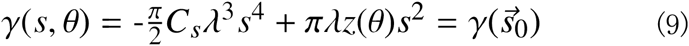

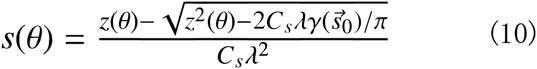

The defocus *z*(*θ*) in Eq. (9) and (10) is calculated using the definition in Eq. (3). Here, I use a positive value for *z*(*θ*) for underfocus. The correct solution for *s*(*θ*) derived from Eq. (9) is the one that lies within the normal range of frequency (smaller than Nyquist).

### 2.9 Self-consistency verification and micrograph quality evaluation

Most currently available programs, such as CTFFIND3, output a diagnosis power spectrum file that needs to be checked manually. Gctf provides the additional option of automatic checking the results. I check the CTF determination by the self-consistency between global fitting and individual frequency shell fitting (Figure 5). As described above (Section 2.4), for higher frequency the SNR is lower but at the same time a tighter criterion is required for accurate CTF determination. The fitting will become more and more unstable for higher frequency shells. For each shell Gctf recalculates the CTF parameters starting with the global values plus some deliberately added error. The added error is as large as the 90° criterion accuracy for the highest resolution of that shell. In good cases, the refinement will converge to original value, while in bad cases the refinement becomes unstable and eventually diverges. If all frequency shells show perfect convergence, there is almost no doubt that the fitting is correct. If only higher frequency shells fail to converge, the micrograph may still contain useable information up to a certain resolution. In some cases there may be problems with the fitting in all shells or an abnormal signal in a certain frequency shell. Gctf regards these micrographs as unusable.

**Figure 5.**
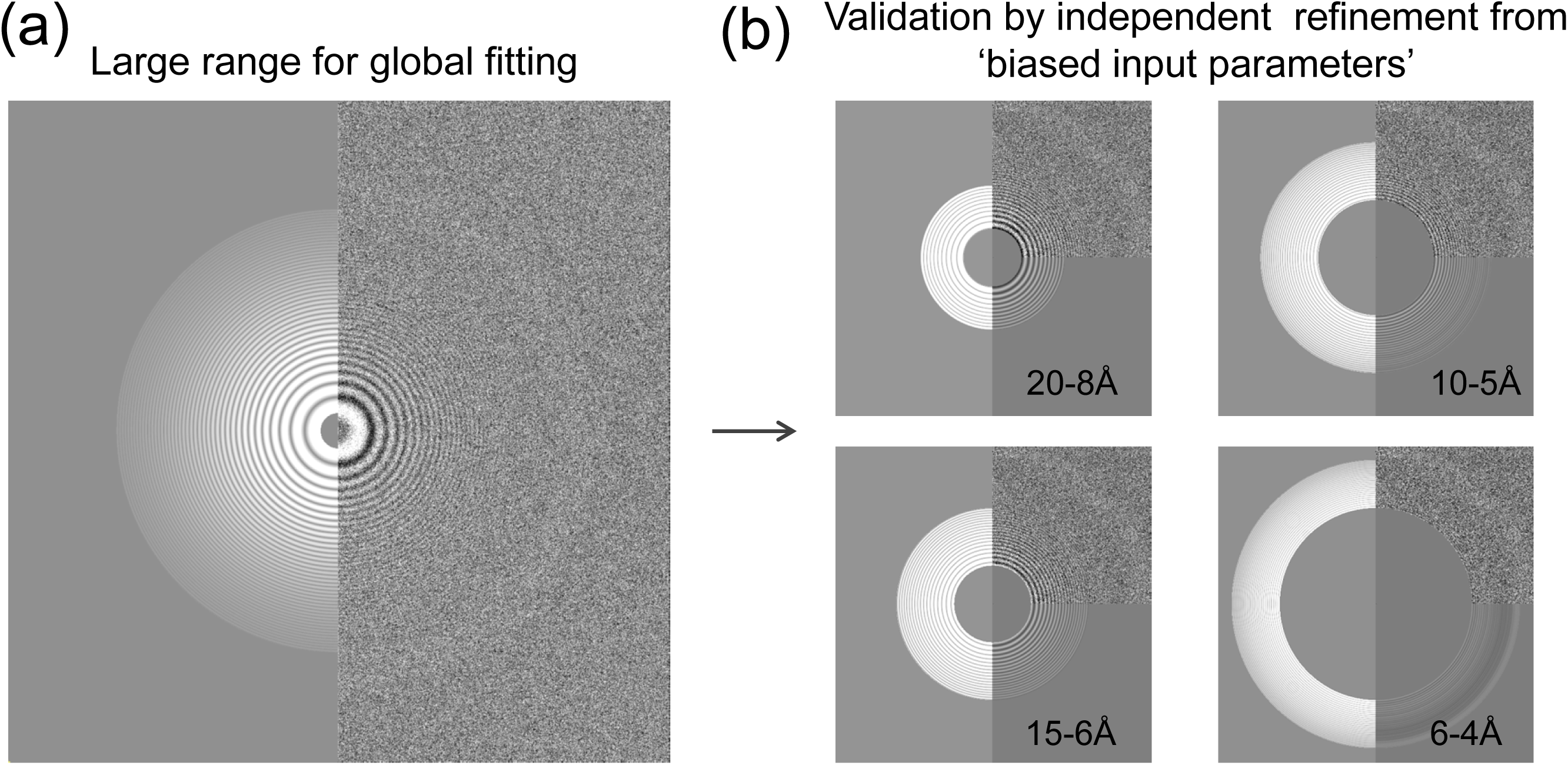
Self-consistency verification of CTF determination and micrograph quality. (a) A typical output power spectrum image for diagnosis; simulation on the left and original on the right. (b) Validation by independent refinement from ‘biased input parameters’ at different resolution shells. The deliberately introduced bias is the same as 90° phase shift criterion for the higher resolution in a certain shell. For example, for the 15-6Å shell, the 90°criterion for 6Å is 914Å; this value is added to the input for refinement.

Gctf also determines the quality of information at different resolution for each micrograph by calculating CCC between the simulated and observed power spectra in each group of continuous Thon rings (6 complete rings form one group in Gctf). For each resolution shell, if the CCC is larger than zero, it is regarded as usable. When the CCC values begin to oscillate above and below zero, the shell is assumed not to contain usable information.

The values determined by Gctf can also be automatically checked for self-consistency in real time according to the historical refinement results. One criterion is that the astigmatism is fixed for a certain dataset, or at least for a certain period of time over which all the parameters of the microscope are relatively stable. Another useful check follows the observation that people tend to collect data at a certain range of defocus for a specific cryoEM sample. Based on these two criteria, the reliability of the defocus from a certain micrograph can be well estimated. If the difference of defocus value from the average is suddenly much larger (e.g. 3 times) than the standard deviation or if the astigmatism suddenly varies more than expected, the micrograph or the CTF determination is potentially abnormal.

### 2.10 Acceleration by GPU, ‘1S2R’ and optimized programming strategy

Gctf was written in the GPU programming language CUDA (C-language version) (https://developer.nvidia.com/cuda-zone). The speed of current high-end GPUs is around several TFLOPS(e.g. NVIDIA GeForce GTX 980 at 4.6 TFLOPS), while high-end CPU is normally ˜100 GFLOPS(e.g. Intel Xeon E5-2643 v2 at 168 GFLOPS). Therefore, programs can be accelerated by tens of times faster using high-end GPU.

In addition to GPU, the fast‘1S2R’ procedure can accelerate Gctf by tens of times more. The acceleration by ‘1S2R’ becomes even more significant than GPU when the step size of defocus used for initial search becomes smaller. This is because Gctf only uses the step size for 1D search rather than 2D. The time of 1D search is actually ignorable compared to the time by other steps (details to be discussed in Section 3.3). In contrast, searching initial defocus is the limited step in ‘2S2R’ procedure as used in CTFFIND3.

The program was further optimized to run as fast as possible by improving the overall strategy(Figure 6). First, instead of sequentially processing each file (Figure 6a), Gctf tries to process an entire dataset containing hundreds or thousands of micrographs together (Figure 6b). This speeds up the program because a lot of computing resources can be shared among the processing of all the files, the hardware only requires initializing once and sharable parameters, values etc. are also only calculated once (Figure 6b). This concept was not only used in the overall processing of hundreds or thousands of micrographs, but also in lots of the sub-procedures in the entire program. Optimization of file reading strategy can also make the speed of Gctf even faster(Figure 6c).

**Figure 6.**
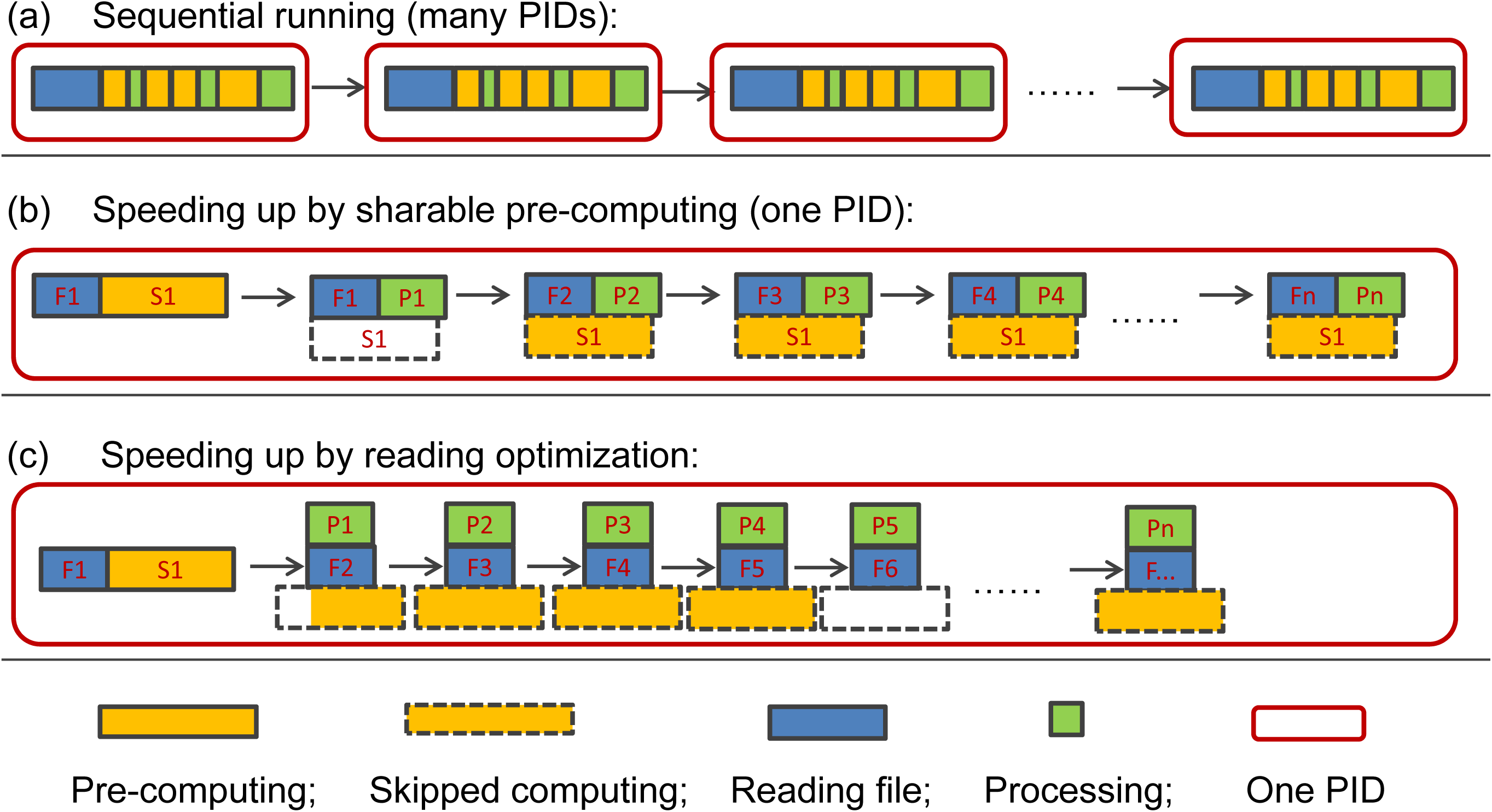
Diagrams comparing speed of runs using different strategies. (a) A traditional strategy for CTF determination. The program processes one micrograph at a time. All micrographs are treated completely independently. Batch processing is normally realized by an external script. (b) An improved strategy to speed up batch processing using a single PID(Process Identifier). In this strategy, a lot of memory, many data and parameters are shareable; some of the frequently used values can be pre-calculated, thus saving significant computing time. (c) A further decrease in run time results from optimization of file reading. Instead of reading one file and then processing it, Gctf can read the next file while simultaneously processing the current micrograph.

Apart from the internal acceleration, I also supply a convenient script to help users take advantage of multiple GPU resources in their local area network. This can almost linearly speed up the program on multiple nodes/workstations/PCs using fast parallel file systems.

### 2.11 Typical input/output of Gctf

The program processes multiple micrographs in batch mode without the need for additional scripting. The basic output of Gctf contains four parts: 1) the standard output which gives real time information about the processing of the micrographs; 2) the log files which contain all the necessary input and output parameters for each micrograph; 3) the diagnosis power spectrum file in MRC format; 4) a STAR file which contains all the determined CTF parameters. Local or movie refinement outputs additional files. The program is fully compatible with Relion and the log files and output STAR files can be directly used for further 2D or 3D classification. The STAR files can either contain CTF parameters for entire micrographs or individual particles.

CTF parameters from the text log file can also be easily extracted and used for other programs such as EMAN (Ludtke et al., 1999), Spider (Shaikh et al., 2008) and Xmipp (Sorzano et al., 2004) et al. The diagnosis file is similar to that of that of CTFFIND3 but contains additional user defined options. Also, Gctf benefits from applying a B-factor to make the contrast between simulated and observed Thon rings more comparable at high frequency.

### 2.12 Variable types of application by Gctf

The basic usage of Gctf is CTF determination of the whole micrograph. It can also be used to refine CTF parameters provided by users. In addition, it is possible to do CTF determination and refinement for particle stacks if original micrographs are not available in some cases. More advanced aspects such as local and movie CTF refinement are also available. Local refinement can be applied to significantly improve the defocus of each particle in a micrograph. This has improved resolution of several real 3D reconstructions (section 3.6). Movie CTF refinement can be used to determine defocus changes with time or dose (section 3.5). Self-consistency verification or micrograph quality evaluation during CTF determination is optional to users. The option can be used to automatically check CTF determination results and report the overall quality of each micrograph. It is also optional to users to flip the phases automatically after CTF determination. All these kinds of processing only require a single and simple command, running in batch mode for the entire dataset instead of only one micrograph.

## 3. Results and discussion

### 3.1 Datasets used for testing Gctf

Supplemental Table 1 shows all the tested datasets presented in this paper. The following three datasets were used in multiple situations. One micrograph of an empty carbon grid with a large astigmatism was used to test the overall procedure of Gctf. A cryoEM dataset of dynactin on thin carbon film was used to test speed, local refinement and movie refinement. A cryoEM dataset of *HAV* in pure ice with a relatively large astigmatism was used to test the EFA method and local refinement.

### 3.2 Stability and reliability of modified ‘1S2R’ procedure

To test the stability of each step in the methods used in Gctf, I first plotted the cross-correlation functions(CCF) of the 1D averaged power spectrum and simulated CTF curves over the a range of defocus values for several micrographs (Supplemental Figure 1). I tested micrographs collected under a variety of conditions, including different doses, detectors, magnifications, types of grid, and levels of astigmatism. The CCF plots each showed a single clear peak indicating that it was possible to unambiguously determine the averaged defocus (Supplemental Figure 1).

One of the major concerns for estimating the averaged defocus using rotational averaging of the power spectrum is that it might fail due to large astigmatism. However, I tested a series of micrographs with a range of astigmatisms (Supplemental Figure 1) and I found that even in the case of an astigmatism of 10000Å Gctf was still able to identify the correct defocus (dataset 1, Figure 7a). Considering the astigmatism of micrographs under normal cryoEM imaging conditions is less than 1000Å, and at most 2000Å, I conclude that an initial 1D search works for almost all normal cryoEM micrographs. For some more challenging cases, Gctf provides additional options(REBS) to deal with them (Section 2.7 and 3.6).

**Figure 7.**
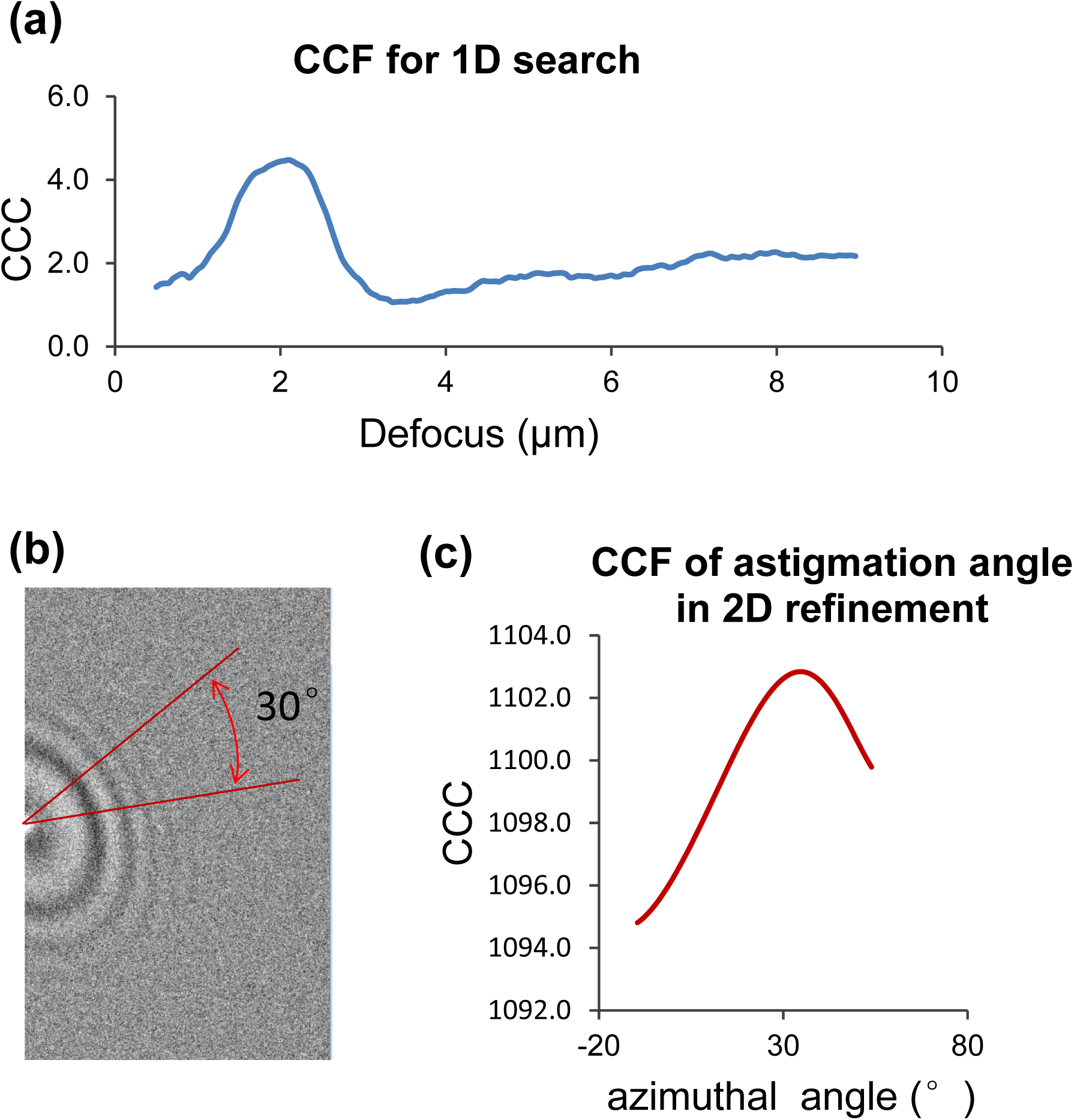
Examination of the ‘1S2R’ strategy in Gctf. (a) The cross-correlation function of 1D search. A single peak is clearly distinguished from the rest, indicating that the determination of averaged defocus is reliable even if the astigmatism is significant. (b) A half image of the power spectrum is divided into several parts and the azimuthal angle is assigned by large step search within a 30° wedge. (c) Cross-correlation function over azimuthal angle in 2D refinement is plotted, showing a clear single peak near the true value. Cross-correlations are not normalized for 1D search and 2D refinement in order to allow a higher speed of GPU programming, but the final output will be normalized in Gctf.

Since a 2D digital micrograph normally contains thousands of times more elements than a 1D curve, an exhaustive search for the defocus and astigmatism would be incredibly slow in 2D. On the other hand, directly refining the three parameters of *z*(***u***, ***v***, ***θ***) in 2D starting from the results of a 1D search has the problem that it can fall into a local minima. To best balance the accuracy of a global search and the speed of a Simplex refinement, Gctf does an initial 2D search based on the 1D result using several large steps to localize the azimuthal angle *θ* within a 30° wedge under the constraints from the relationship between ***z*_*u*_** and ***z*_*v*_** (Figure 7b). Next it does a 2D refinement for all three parameters of *z*(***u***, ***v***, ***θ***). Tests on many datasets show starting from this value of *θ* allows for a robust refinement of *z*(***u***, ***v***, ***θ***) by the Simplex method (Supplemental Figure 1).

### 3.3 Speed test and comparison

I tested the speed of Gctf using different parameters on different devices. In general the speed can be comparable to that of simply reading the files. The kernel of CTF fitting only takes ˜0.1 second, using a currently available high-end GPU (i.e. Nvidia GTX 980). In addition to GPU acceleration, which has been shown to accelerate many programs by tens of times(Li et al., 2010; Xu et al., 2010; Zhao and Chu, 2014), Gctf also has significantly improved algorithms and programming strategy (Figure 2, 6 and 8a). A detailed speed comparison with CTFFIND3 was done (Figure 8b). In general, the speed of CTFFIND3 is limited by both the step size of the defocus search and the FFT box size, whereas the speed of Gctf is independent of either. The reasons are different for these two parameters. Gctf uses the fast procedure ‘1S2R’ while CTFFIND3 use the procedure ‘2S2R’. The step size of defocus search in Gctf is only used in 1D and does not affect the speed of 2D refinement by Simplex method. Since the 1D search takes less than 0.0001s, Gctf is significantly accelerated by this ‘1S2R’ strategy. The FFT is performed on highly-parallel GPU and the speed does not decrease with the increase of box size until it is larger than 1024. For typical usage (box size 512 pixels and search step 500Å), Gctf can process more than 1000 micrographs in the time CTFFIND3 requires to process one. However, Gctf uses a box size 1024 by default for better diagnosis of high defocus micrographs at high frequency. Using this box size the acceleration is even more significant.

**Figure 8.**
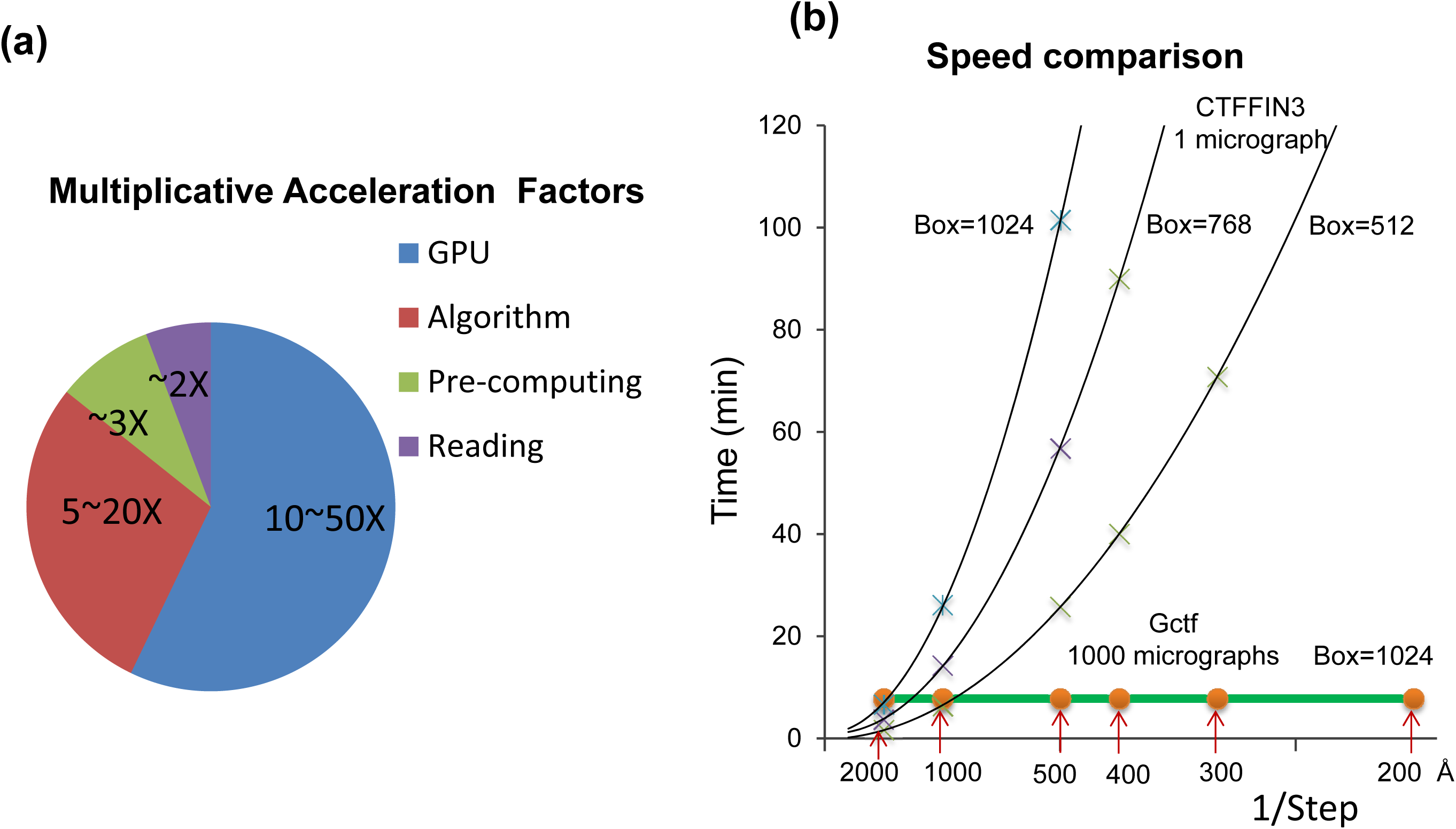
Acceleration in Gctf and comparison with CTFFIND3. (a) A diagram showing the accelerating factors Gctf tries to optimize. One significant factor is GPU, which can easily speed up tens of times. The optimized procedure ‘1S2R’ contributes to several tens of times acceleration. This becomes more significant when the step size is smaller or the box size is larger. Pre-computing and reading optimization, although not as significant GPU and algorithm acceleration, can make the speed 4˜5 times faster together. Notably, the effect of these accelerating factors is multiplicative rather than additive. Therefore, Gctf is much faster when they are combined. (b) Speed comparison between CTFFIND3 and Gctf. The box sizes for FFT are marked on the side of each speed curve; x-axis represents the defocus step size for search in reciprocal space(larger value represents smaller step), the real value of the step size are marked by the red arrow. CTFFIND3 was running on one of the latest high performance CPU, Intel Core(tm) i7-4790K @4.00GHz; Gctf was running on the same computer using the GeForce GTX980 GPU.

Gctf is capable of handling different types of CTF determination and refinement. The limiting factor is mainly the file reading or network speed (Table 1). In a test using dynactin micrographs (dataset-5, Supplementary Table 1), the average speed of Gctf can be even accelerated 3 times (from 0.75s to 0.26s) simply by using a fast SSD disk. This indicates that the speed limitation is at the file reading step. For movie CTF refinement reading takes 5-30s but once the movie is read into GPU RAM, processing takes less than 1s. The limitation for local refinement, however, is mainly the particle number which is approximately linear to the fitting time. It’s much slower than global CTF determination, but very useful to improve the CTF parameters of each particle (Section 2.5 and 3.7).

**Table 1.** Typical speed of Gctf for different types of application^a^

|  | Dataset-3 | Dataset-9, | Dataset-5 , | Dataset-6,<br>(local<sup>b</sup>) | Dataset-5,<br>(movie<sup>c</sup>) |
| --- | --- | --- | --- | --- | --- |
| GTX750 Ti, HDD | 0.29s | 0.59s | 1.14s | 4.46s | 4.27s |
| Tesla K40, NFS | 0.65s | 0.76s | 1.00s | 3.58s | 7.88s |
| K5000, NFS | 0.68s | 1.18s | 1.34s | 4.31s | 13.07s |
| GTX980, NFS | 0.36s | 0.60s | 0.75s | 3.48s | 5.60s |
| GTX980, SSD | 0.10s | 0.16s | 0.26s | 1.93s | 2.20s |

a: Time of each micrograph in average
b: 44 particles in total
c: 20 frames, coherent averaging
HDD: Serial ATA Hard Disk Drive
NFS: Network File System,
SSD: Solid State Disk

### 3.4 Self-consistency of CTF determination and micrograph quality

I estimated the convergence and accuracy of the CTF determined by Gctf using the described method above (Section 2.9). The estimation of accuracy depends on the bias and error of fitting. They can both affect the final quality of CTF determination but for different reasons. Bias comes from over fitting of strong false signals in a certain range of frequency, which normally derives from the background (e.g. big ice contamination) or significant structural information (e.g. ring-like structure or ring-like features in the structure). Gctf deals with bias by independent resolution shell refinement and uses these results to estimate the accuracy and reliability of CTF determination (Section 2.9). In contrast, random error mainly reflects the quality of the micrograph itself and only affects quality of CTF determination at high frequency. It is affected by many experimental factors: ice thickness, alignment of the microscope, detector DQE and so on. Based on the estimated CTF fitting bias and error, Gctf will output an overall diagnostic check of self-consistency for each micrograph. It uses a score representing different levels of quality up to a certain resolution: 0=wrong, 1=bad, 2=usable, 3=good, 4=very good and 5=perfect.

A summary of CTF determination results by Gctf is presented in Supplementary Table 2. Note that the CTF determination results which were “BAD” or “WRONG” could always be traced to a problem of the micrographs. Several representative CTF determination results that were evaluated as “BAD” or “WRONG” were shown in Supplementary Figure 2. In general, the quality of micrographs on direct electron detectors is much better than those on CCD. In the case of a high quality dynactin dataset on Falcon II detector, 99.7% or 97.9% micrographs are evaluated as usable based on 8Å or 4Å criterion. The HAV dataset on K2 summit detector shows comparable results. The chaperonin dataset(Zhang et al., 2013) on UltraScan4000 CCD shows worse results, 92.7% are evaluated as usable based on 8Å criterion and only a quarter is evaluated as usable based on 4Å criterion.

Gctf also provides a power spectrum file for manual diagnosis. In addition to the observed and simulated power spectra, the user can also view a rotational average and/or EFA power spectra (Figure 9a). For high-quality micrographs of grids covered with a thin carbon film the observed power spectrum can show Thon rings up to 4Å. But for micrographs of pure ice, the rings usually disappear between 6 to 8Å even after rotational averaging. I recommend using equiphase averaging(EFA), which makes it possible to view Thon rings up to near-atomic resolution for many micrographs taken on direct detectors, or even for individual particles.

**Figure 9.**
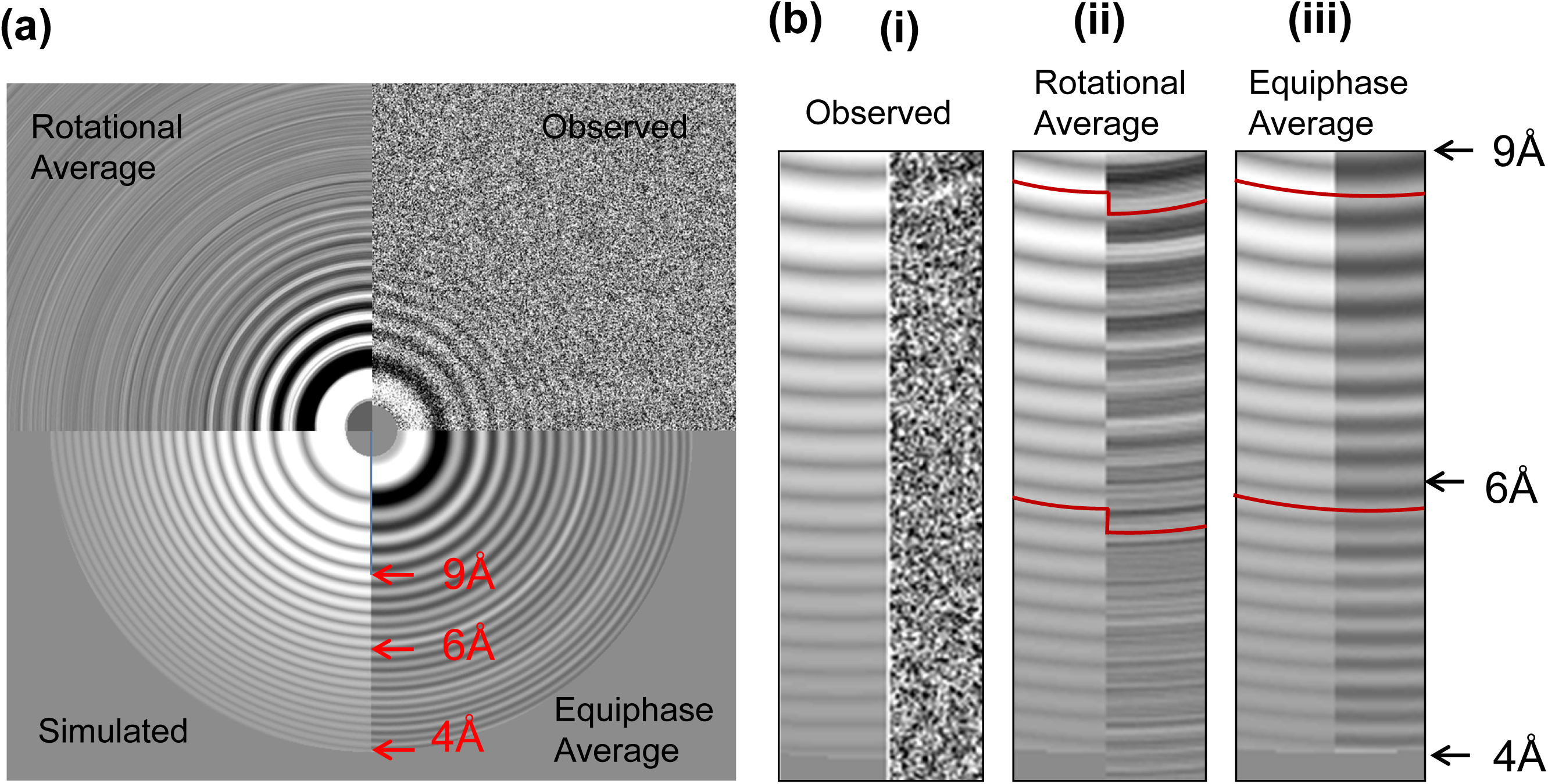
Equiphase average for better diagnosis. (a) Different power spectrum images of a typical cryoEM micrograph of HAV(dataset-8) with ˜1800Å astigmatism; left top: normal power after background reduction; right top: normal rotation average; left bottom: equiphase average; right bottom: simulated. (b)Enlarged region from 9Å to 4Å for detailed comparison of different diagnostic methods. Thong rings on the left sides in all the three images represent the simulated power spectra; the right sides represent the observed power spectra visualized in different methods. Obviously, Thong rings from the original power spectrum (i) are almost invisible at higher than 9Å even after background reduction. After rotational average (ii), the rings become clearer but significantly off the correct peak because of the large asgtigmation. Some rings are almost in reverse contrast, indicating the rotational average is meaningless at such resolution. The equiphase average (iii) makes all the rings clearly visible up to 4Å.

### 3.5 Robustness

A good way to test the robustness of a CTF determination program is its tolerance for dealing with different ranges of input parameters. I used Gctf to determine the CTFs of different datasets from three examples, dynactin (in dataset-6), HAV (in dataset-9) and a pure carbon on Quantifoild grid (in dataset-7) using variable ranges of four parameters: resolution, astigmatism, box size and B-factors.

I first tested the ability of Gctf to handle different specified resolution ranges. It could correctly determine the defocus of an HAV dataset in pure ice with high resolution cutoffs between Nyquist and 20Å. Likewise all the CTF determination results were very similar for a low resolution cutoff between +∞ to ˜8Å when high resolution cutoff was fixed at 3Å (Figure 10a, Movie S1-S3). When this cutoff was smaller than 8Å, CTF determination became unstable and the results are not reliable. However, it’s encouraging that most of the results were still near the correct values for low resolution range from ˜8Å to 4Å. Even if very narrow ranges of resolution around 4-3Å were used, the errors were only ˜400Å. On the other hand, Thon rings were visible only to ˜8Å resolution by eye without using EFA. For dynactin, which was collected on thin carbon film, the resolution cutoff range was more robust (Supplemental Figure 3a). This means the default resolution range (50-4Å) for Gctf does not normally need optimization. This is quite helpful for automatic cryoEM data processing.

**Figure 10.**
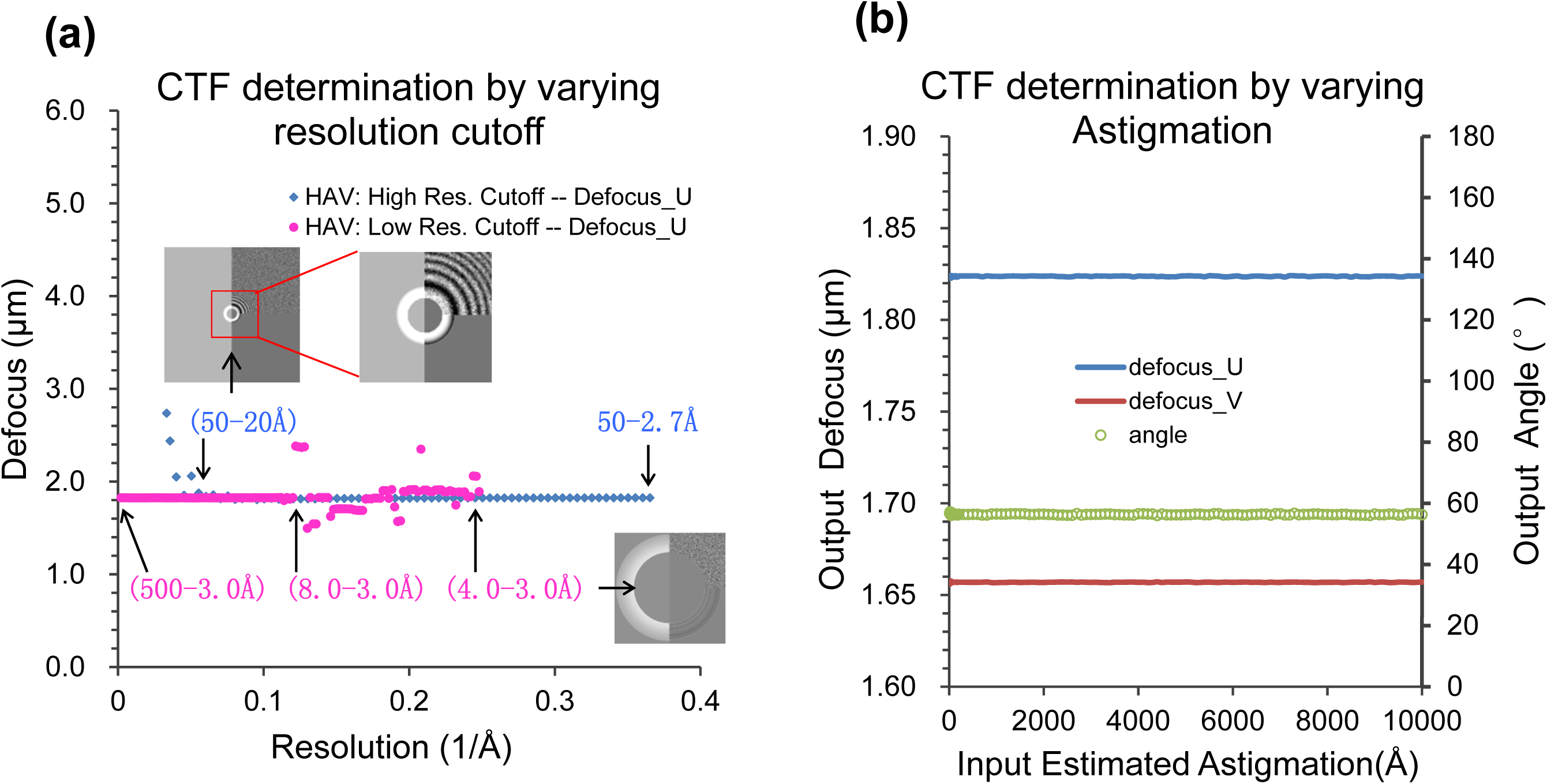
Robustness test of CTF determination using varying resolution cutoff or estimated astigmatism as input. (a)Typical CryoEM micrograph of HAV in pure ice was selected and systemically examined to test the robustness of Gctf. The stable range for the high resolution cutoff(blue) is from ˜20Å to Nyquist; The stable range for lowest resolution cutoff is +∞ to 8Å. The blue points represent results by high resolution cutoff and the red points by low resolution cutoff. (b) The input values of astigmatism ranging from 10Å to 10000Å were used as initial estimation for CTF determination. All input values in this range generated almost identical results. Therefore, there is no need for optimizing the input astigmatism. Blue and red lines represent defocus U and V respectively. Green line represents the azimuthal angle.

I also tested the range of astigmatism input values that Gctf can tolerate (Figure 10b). It can find the correct astigmatism with given starting values between 10Å to 10000 Å for an actual astigmatism of ˜1800Å. This suggests that Gctf can refine the astigmatism of any practical cryoEM micrographs, which usually ranges from 200 Å to 2000Å, using the default parameter (1000Å).

I next tested how the Bfactor could affect the CTF determination in Gctf. The relatively stable range is around 0˜to 1500 for the HAV dataset(Supplemental Figure 3b). Using the default Bfactor values(150Å^2^), Gctf is able to accurately determinate the CTF for this dataset. In some challenging cases(e.g. very big astigmation), the optimization of Bfactor might help to get better accuracy.

I deliberately collected several micrographs with larger astigmatism on carbon film as a case study (in dataset-7, Supplemental Table 1). The averaged defocus determination on one of the micrographs was always correct, but the astigmatism estimation could easily fail using default settings in Gctf (Supplemental Figure 4a). I provide two approaches, Resolution Extension(RE) or Bfactor Switch(BS) or the combination(REBS) to deal with similar cases as previously described (Section 2.7). The azimuthal angle can be accurately determined using either of the following parameters: (1) 50-10Å resolution range, Bfactor=150Å^2^; (2) 50-4Å resolution range, Bfactor=500Å^2^. However, there is 200Å˜ 300Å defocus error using either of these options. Based on either of the two results, a second step of CTF refinement using 15-3Å resolution range and Bfactor=50Å^2^ gives very accurate results as shown in Supplemental Figure 4 using EFA. Comparable results could not be obtained by a single step of CTF determination without using the REBS procedure in this case.

The selection of box size might affect the accuracy of CTF determination. In general, larger box size is better than smaller. This is because the oscillation of CTF is severe at high frequency. If the box size is too small(e.g. 128 or 256), the sampling ratio at high frequency is not sufficient to separate adjacent Thon rings in the power spectra for high defocus micrographs. This will affect both CTF determination accuracy and manual diagnosis(Supplemental Figure 5). The big error (˜1100Å) due to very small box size (128) is not acceptable for near atomic reconstruction according to the criteria in Figure 3. On the other hand, the box size should not be too big for local CTF refinement. Otherwise, the improvement by local CTF refinement is not significant. By default, Gctf uses box size 1024 for global CTF determination and 512 for local CTF refinement.

### 3.6 Challenging cases of CTF determination

For easy cases, i.e. micrographs with carbon film at high dose, results from all available CTF determination programs are comparable. The exceptions are cases with large astigmatism since some programs, only determine averaged defocus. The differences among programs become significant for challenging cases. I have shown that Gctf can accurately determine the CTF for several challenging cases, i.e. low contrast micrographs collected on CCD (Supplementary Figure 6a), micrographs with smaller defocus(Supplementary Figure 6b), very large defocus(Supplementary Figure 6c) or very large astigmatism(Supplementary Figure 4) and samples containing ring-like features(e.g. DNA origami), single frames from a movie (Figure 11a) and so on. Especially, the power of Gctf is demonstrated by its ability to determine the CTF of single frames from a movie with doses of only 1-2 e/Å^2^ (Figure 11a, Movie S4). By averaging adjacent frames (e.g. 5-10), the results are accurate enough to detect the changes of Z-height (Figure 11b).

**Figure 11.**
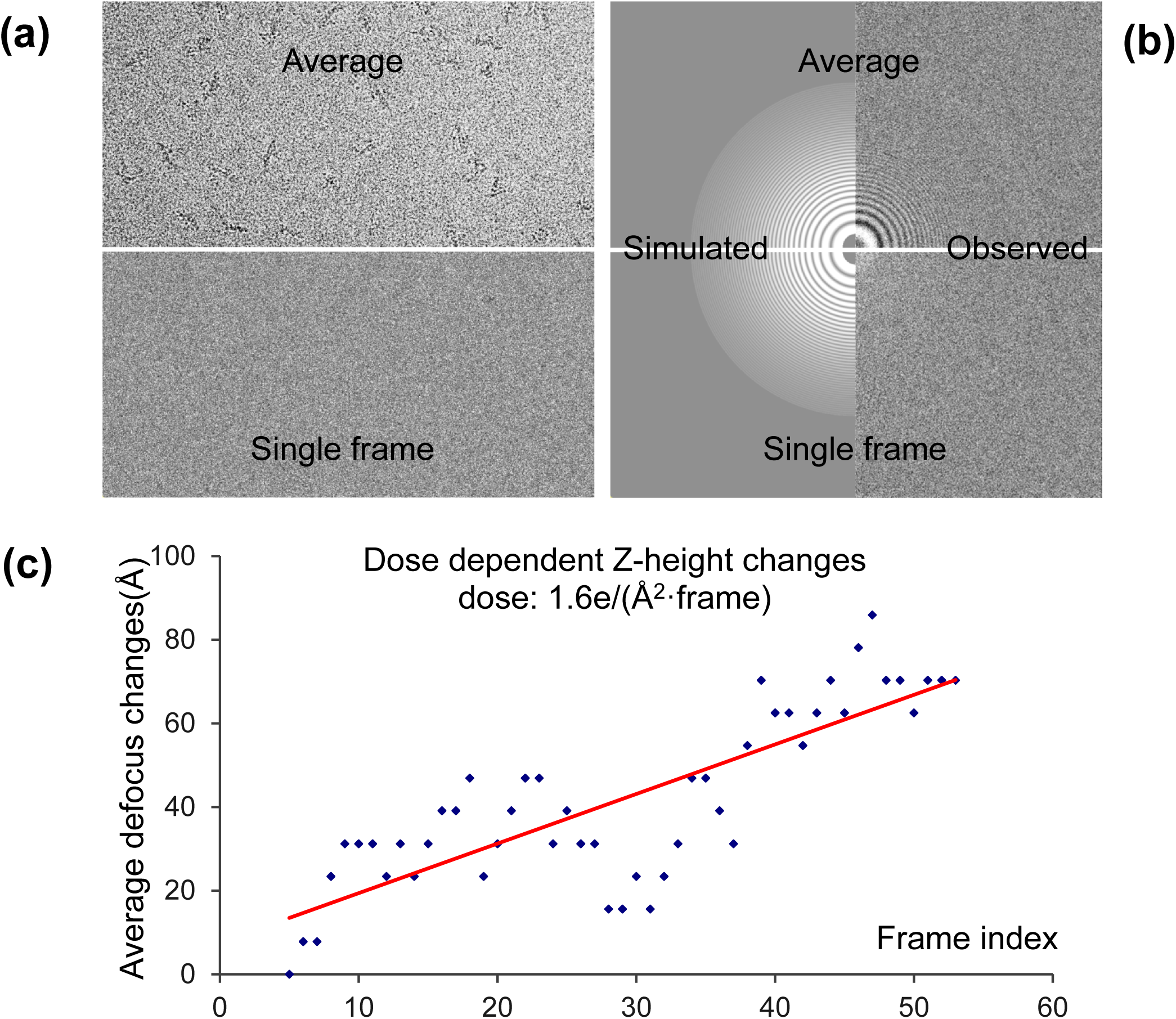
CTF determination of single frames of a movie. (a)Averaged movie (top) and single frame(bottom) of dynactin(dataset-5). Movie was taken on FEI Titan Krios, Falcon II detector at the dose of 1.6e/(Å2·frame). (b)CTF determination using the averaged movie(top) or a single frame(bottom). For both images, the left is simulated CTF and the right is observed power spetrum. The determined defocus *z*(*u*,*v*,*θ*) of the averaged movie is (41642.58Å, 41140.62Å, 61.67°) and the first frame is (41711.86Å, 41196.36Å, 52.80°). The difference is (69.28Å, 55.74Å, 8.87°). (c) The changes of averaged defocus (*z*_*u*_ + *z*_*v*_)/2 with the accumulation of doses on the micrograph. Slightly different from (b), the CTF determination for each frame was performed by averaging 9 adjacent frames(e.g. 11-19 for frame 15) to enhance the SNR.

### 3.7 Significant improvement of defocus accuracy by local refinement for each particle

Gctf can estimate the defocus for each particle accurately using the current method described in Section 2.5. One good example, from a dynactin micrograph with a typical defocus variation is shown in Figure 12. Since the cryoEM grid was coated with thin carbon film and the quality of this micrograph is very good, the peaks of the rotational averaged Thon rings are clearly visible for each particle up to 4Å. A comparison of two representative particles shows that the position of their Thon rings is obviously shifted. The peaks in the power spectrum are almost reversed at higher than 5Å resolution. A clear comparison between these two particles is shown in Movie S5.

**Figure 12.**
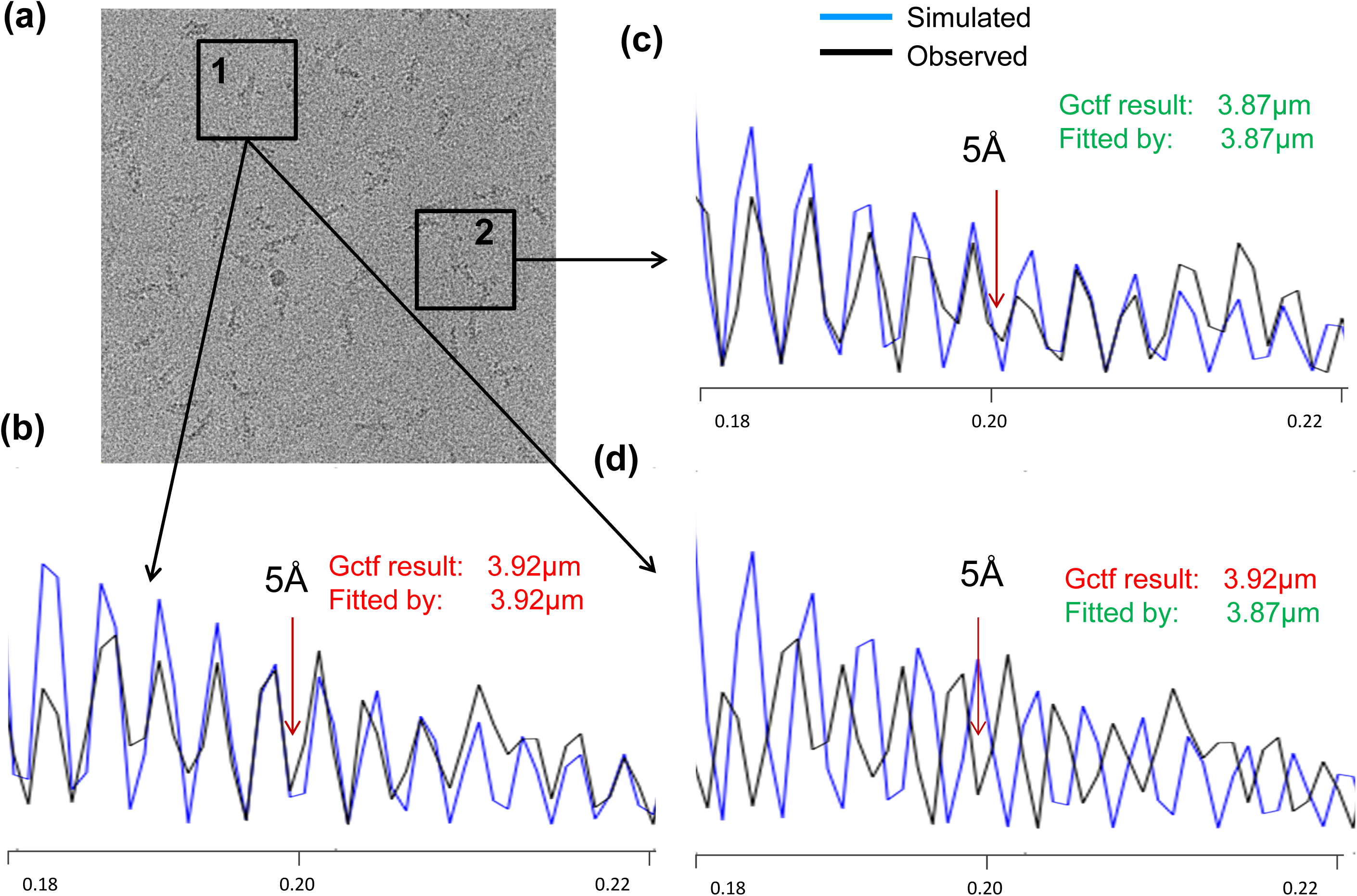
An example showing the importance of local CTF refinement. (a) The raw micrograph of dynactin(dataset-6). (b) The local defocus for this particle determined by Gctf is 3.92µm. The red arrow indicates the fitting is almost perfect up to 5Å. The black curve represents the rotational average of the power spectrum of particle 1; the blue curve represents the comparison with Gctf determination. (c) Similar to (b) but for particle 2. The defocus of this particle is 3.87µm. The fitting of CTF is also perfect up to 5Å. (d) Comparison between the observed power spectrum of particle 1 and simulated power spectrum using defocus of particle 2. In contrast to or (c), the simulated CTF curve does not fit the observed curve at high resolution, indicating the importance the local refinement for near atomic resolution reconstruction.

I carefully examined many micrographs and found the local defocus variation can be much larger than expected from tilt micrographs. I plotted the maximum and standard deviation of the averaged local defocus for all micrographs in dataset-6 (Supplementary Figure 7). The standard deviations of local defocus for over 50% micrographs are actually larger than the theoretical value of a micrograph with 10 degree tilt. The maximum local defocus deviations for about one third micrographs are even larger than the theoretical value of a micrograph with 15 degree tilt. On the other hand, the grid was proved to be flat by the local defocus variation in several regions of completely burned carbon(Supplementary Figure 8a). In these regions the local defocus variation is smaller than 30Å which is theoretically equivalent to ˜3 degree tilt. Therefore, I conclude the local defocus variation in cryoEM micrographs is not attributed to tilt, but other factors such as uneven ice, carbon support or charging.

In addition to self-consistency verification of global CTF determination, local defocus variation determined by Gctf can also be used to detect abnormal micrographs. Micrographs with very big local defocus deviation(e.g. maximum deviation >1000Å) were always low-quality(Supplementary Figure 8b) or partially unusable(Supplementary Figure 8c,d).

The speed and accuracy of Gctf was helpful during the determination of the near-atomic resolution cryoEM structure of dynactin (Urnavicius et al., 2015) (Figure 13a) and HAV (Figure 13b, by courtesy of X. Wang). I tested how much local defocus determination affected the final reconstruction of both samples. Generally, the improvement depends on the magnitude of the defocus variation in the micrographs. Local defocus refinement by Gctf never made the final reconstructions worse. In one of the best cases for dynactin in a thin carbon layer support, the resolution was improved from 4.7Å to 4.4Å (Figure 14a). In the case of HAV, which was in pure ice and therefore assumed to be less affected by uneven support, local refinement could also improve the resolution from 3.5Å to 3.4Å (Figure 14b).

**Figure 13.**
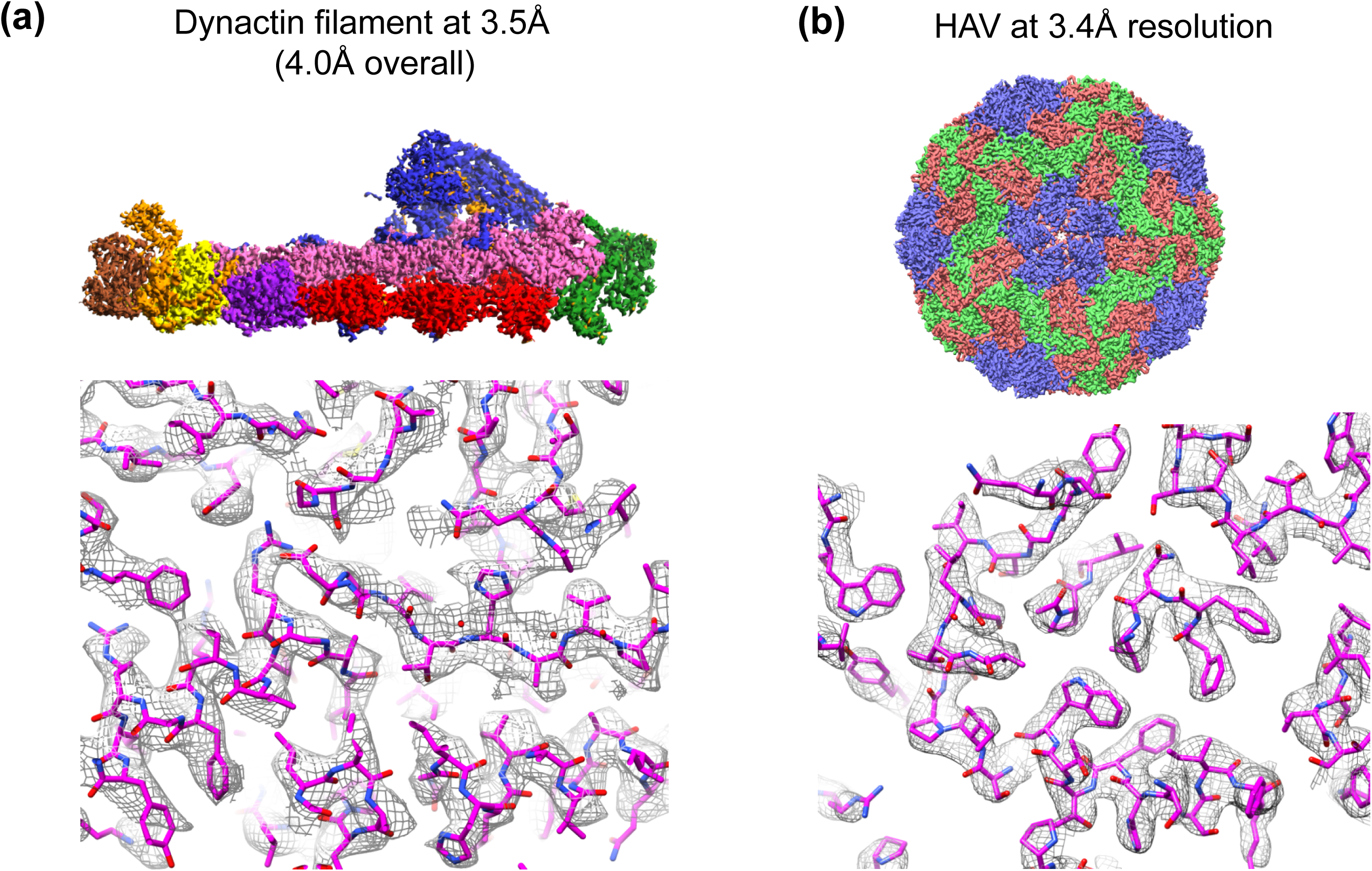
Representative structures that benefits from both the speed and accuracy of Gctf. (a) Dynactin filament at 3.5Å resolution. The whole complex is solved to 4.0Å resolution overall. This result is a combination of several datasets. (b) HAV at 3.4Å resolution(by courtesy of Dr. X.X Wang).

**Figure 14.**
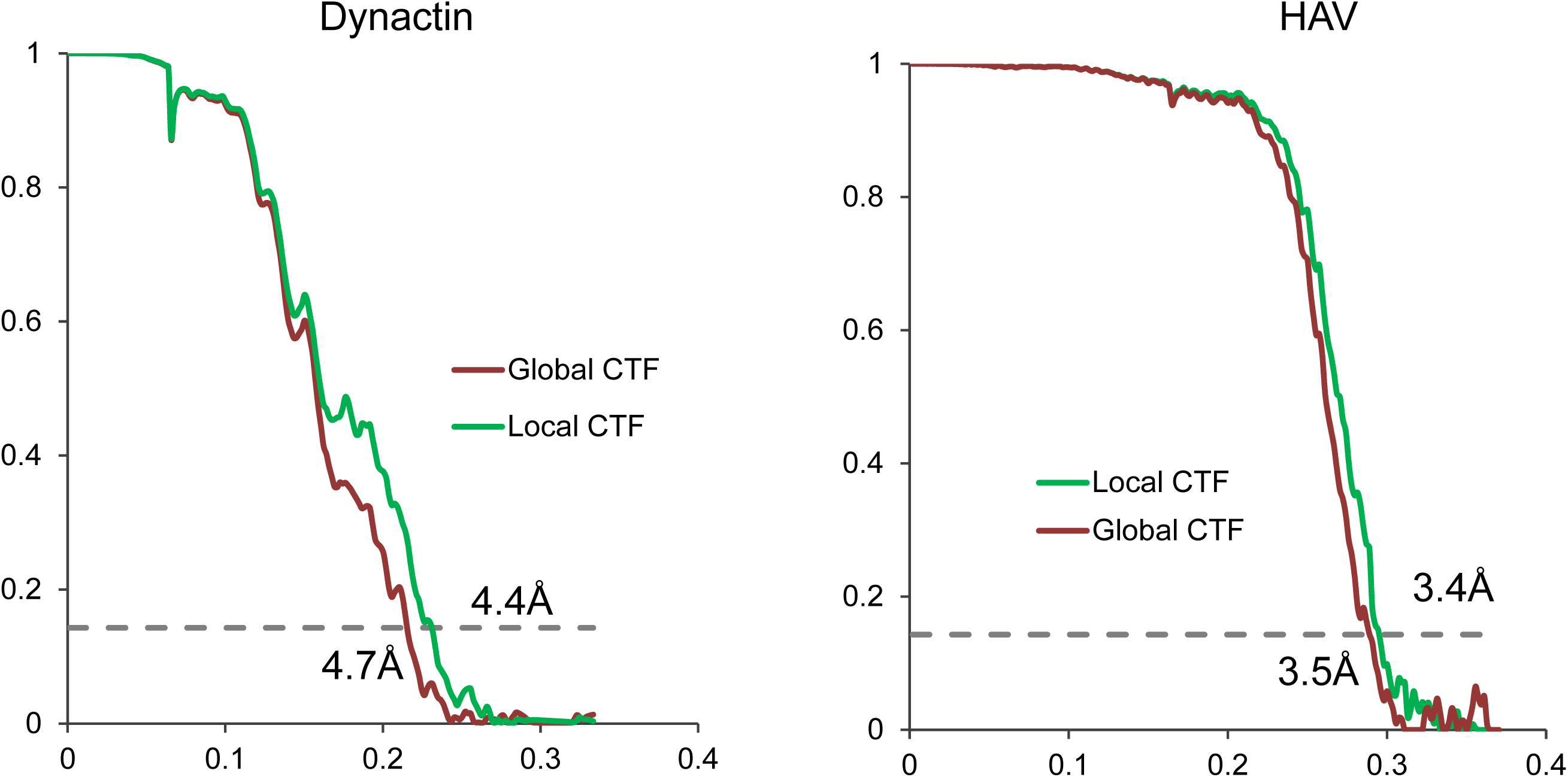
FSC comparison between global and local CTF. (a) Comparison of Dynactin FSC curves with(green) and without(red) doing local defocus refinement. 86916 particles were used for final reconstruction from dataset-7. (b) Comparison of FSC curves of HAV with(green) and without(red) doing local defocus refinement. 2025 particles were used for final reconstruction in dataset-8.

## 4 Conclusion

Gctf is a convenient, accurate, robust and very fast CTF determination and correction program. GPU acceleration, the fast ‘1S2R’ procedure and optimized programing strategy all together have made it a real-time program. Approaches of self-consistency verification and micrograph quality evaluation have also been proposed for automatic CTF determination and micrograph selection. Approaches for local CTF refinement of each particle in a micrograph or frames in a movie have been proposed to improve the accuracy of CTF determination. Extensive practical tests proved its power to facilitate cryoEM image processing and could improve the final resolution of 3D cryoEM reconstructions in some cases.

## Abbreviations

1D (2D, 3D): One(two, three) dimensional
cryoEM: cryogenic electron microscopy
CCC: Cross-correlation Coefficient
CCD: Charge Coupled Device
CCF: Cross-correlation Function
CTF: Contrast Transfer Function
DQE: Detective Quantum Efficiency
EFA: Equiphase Averaging
FFT: Fast Fourier Transform
GPU: Graphic Processing Unit
HAV: Hepatitis A virus
NFS: Network File System
SNR: Singal Noise Ratio
SSD: Solid State Disk

## Appendix

All supplementary tables and figures are available online. During the preparation of this paper, a bench mark study of CTF determination on challenging cases was published (Marabini et al., 2015). I therefore downloaded all the data sets for testing Gctf. All the results are attached for comparison (Supplementary Table 3 and 4).

At the time Gctf when was pre-released in MRC-LMB (Cambridge), STRUBI (Oxford), Institute of Biophysics (Beijing) or by personal communication, another fast program CTFFIND4 (a successor of CTFFIND3) was also released. Therefore, I did additional comparison of speed among CTFFIND3, CTFFIND4 and Gctf(Supplementary Figure 9). CTFFIND4 is ˜10X faster than CTFFIND3, but still using the ‘2S2R’ procedure which is limited by both box size for FFT and step size for searching initial defocus. The speed of Gcf using a single GPU on a workstation is several times faster than CTFFIND4 using 120 CPU cores on a high-performance computer cluster.

## Acknowledgement

This work was funded by the Medical Research Council, UK (MC_UP_A025_1011) and a Wellcome Trust New Investigator Award (WT100387) to Dr Andrew Carter, in whose lab the Gctf program was developed. I thank Dr. Andrew Carter for help with writing the manuscript and Dr. Aristides Diamant for proof reading. I thank Dr. Chris Russo, Dr. Sjors Scheres and Dr. Richard Henderson for their discussion on CTF determination and valuable suggestions on the both the program and this paper. I also thank Dr. Shaoxia Chen, Dr. Christos Savva and Dr. Greg McMullan for their support for electron microscopy; Dr. Jake Grimmett and Dr. Toby Darling for support with scientific computing; Linas Urnavicius for supplying the dynactin complex; Dr. Xiangxi Wang and Ling Zhu for offering their cryoEM micrographs of HAV as a case study; Dr. Fei Sun for offering the cryoEM micrographs of Chaperonin. I also thank all initial users of Gctf, especially Dr. Alan Brown, Dr. Xiangxi Wang, Ling Zhu, Dr. Jun Dong, Shengliu Wang et al, for their valuable feedbacks to further improve Gctf, help on setting up and running Gctf at the MRC-LMB (Cambridge), STRUBI (Oxford), Institute of Biophysics (Beijing) et al.

